# Individual differences in dopamine function underlying the balance between model-based and model-free control

**DOI:** 10.1101/860361

**Authors:** Ying Lee, Lorenz Deserno, Nils B. Kroemer, Shakoor Pooseh, Liane Oehme, Dirk K. Müller, Thomas Goschke, Quentin J.M. Huys, Michael N. Smolka

## Abstract

Reinforcement learning involves a balance between model-free (MF) and model-based (MB) systems. Recent studies suggest that individuals with either pharmacologically enhanced levels of dopamine (DA) or higher baseline levels of DA exhibit more MB control. However, it remains unknown whether such pharmacological effects depend on baseline DA.

Here, we investigated whether effects of L-DOPA on the balance of MB/MF control depend on ventral striatal baseline DA. Sixty participants had two functional magnetic resonance imaging (fMRI) scans while performing a two-stage sequential decision-making task under 150 mg L-DOPA or placebo (counterbalanced), followed by a 4-hour ^18^F-DOPA positron emission tomography (PET) scan (on a separate occasion).

We found an interaction between baseline DA levels and L-DOPA induced changes in MB control. Individuals with higher baseline DA levels showed a greater L-DOPA induced enhancement in MB control. Surprisingly, we found a corresponding drug-by-baseline DA interaction on MF, but not MB learning signals in the ventromedial prefrontal cortex. We did not find a significant interaction between baseline DA levels and L-DOPA effects on MF control or MB/MF balance.

In sum, our findings point to a baseline dependency of L-DOPA effects on differential aspects of MB and MF control. Individual differences in DA washout may be an important moderator of L-DOPA effects. Overall, our findings complement the general notion where higher DA levels is related to a greater reliance on MB control. Although the relationship between phasic DA firing and MF learning is conventionally assumed in the animal literature, the relationship between DA and MF control is not as straightforward and requires further clarification.

## Introduction

Reinforcement learning (RL) has taken a foothold in the study of action control [1]. Two classes of RL methods, namely “model-free” (MF) and “model-based” (MB) learning, have been used to formulate how an agent maximises rewards in uncertain environments [2,3]. While a MF agent reflexively chooses actions that have led to rewards previously, a MB agent plans prospectively using a ‘world model’ and makes choices accordingly [2,4]. The two-stage sequential decision-making task (two-step task, [5]) has been widely used to quantify MF and MB control. In this task, individuals employ a mixture of both MF and MB strategies [5-9].

Cumulative evidence suggests that individuals with higher baseline dopamine levels (baseline DA) exhibit more MB relative to MF control. Supporting this notion, older adults and Parkinson’s disease patients, known to have less overall baseline DA in the brain, exhibit MB decrements while MF control remained intact [10,11]. Further, individuals who have genetically higher DA tone in the prefrontal cortex (PFC) also display higher MB control without MF alterations [12]. A recent F-DOPA PET-fMRI study showed that individuals with higher ventral striatal DA synthesis capacity, indicating higher baseline DA [13], exert more reliance on MB than MF control, show stronger MB learning signals in the lateral prefrontal cortex (PFC) and weaker ventral striatal (VST) MF learning signals [8]. Using a rodent version of the same task, a positive correlation between baseline VST DA and MB control was recently confirmed [14]. Taken together, whereas the relationship to MF control remains unclear, a positive relationship between higher baseline DA and more MB control is supported by a series of human studies and one recent animal study.

In line with this, pharmacological manipulation studies suggest that L-DOPA, a dopamine precursor that increases overall DA in the brain [15], also enhances MB control without altering MF control [11,16]. A prominent hypothesis suggests that drug effects in the dopaminergic system are baseline-dependent and reflect an inverted U-shaped relationship (e.g.[17]). In this regard, it is usually assumed that individuals with low baseline DA levels, in particular in the prefrontal cortex, are sensitive to cognitively enhancing DA drug effects while individuals with high baseline DA levels may even be prone to drug-induced cognitive impairments. Indeed, some studies observed that individuals genetically predisposed to *lower* baseline DA showed greater improvements in learning under L-DOPA than individuals with *higher* baseline DA predispositions [18,19]. Interestingly, drug effects on MB control also seem to depend on other inter-individual differences, in particular in working memory capacity (WMC). For instance, individuals with higher WMC showed greater L-DOPA induced MB enhancement than individuals with lower WMC [11,20]. WMC has been positively associated with striatal DA synthesis capacity [21], which indicates a likely moderation of L-DOPA effects on MB control by baseline DA levels. Hence, the moderation of L-DOPA effects by WMC seems to follow a U-shape relation with no effect on MB control in low DA individuals and enhancement MB control in high baseline DA individuals. To our knowledge, only one animal study has directly investigated the relationship between L-DOPA metabolism measured with ^18^F-DOPA PET and DA changes with L-DOPA and found that increases in DA metabolites elicited by L-DOPA is greater when DA synthesis capacity is higher [22]. Thus, individuals with higher DA synthesis capacity may produce more DA from L-DOPA, thereby enhancing MB control as observed previously in higher-WMC individuals [11].

Based on this background, the relationship between baseline DA and L-DOPA effects on MB and MF control remains to be clarified. Therefore, we scanned participants using fMRI when they conducted a two-step task on two separate occasions, once after L-DOPA administration and another under placebo. They were also subjected to a four-hour ^18^F-DOPA PET scan, a much longer protocol than typically used for measuring DA synthesis capacity, which enables us to derive measures related to later stages of L-DOPA metabolism such as DA washout rate. The ratio of DA synthesis capacity to washout rate, otherwise known as effective distribution volume ratio (EDVR), indicates ^18^F-DOPA trapped in the striatum at *steady-state* [23,24]. As such, it could be a more sensitive marker of baseline DA than DA synthesis capacity or DA washout rate alone, as suggested previously for staging Parkinson’s disease states [25]. With this information, we asked whether higher baseline DA levels and L-DOPA induced DA enhancement are associated with a shift of the MB/MF balance towards MB control, and we addressed the central question of this study: is the effect of L-DOPA on the MB/MF balance dependent on baseline DA levels.

## Materials and methods

### Participants

This dataset was collected for investigating dopamine modulation of cognitive control (https://tu-dresden.de/bereichsuebergreifendes/sfb940/research/b-modulatoren/b3). A part of this dataset has been reported previously [20,26]. Of the participants who agreed to participate in the fMRI and PET visits (n = 103), 43 were excluded from the PET visit as they experienced adverse events (e.g. vomiting) during the pharmacological fMRI experiment (n = 13), had poor data quality (e.g. excessive max. translation > 3mm from reference image, missing trials > 25%) (n = 9), incidental findings (e.g. brain atrophy) (n = 1), incomplete data due to technical issues (e.g. data lost, drug manipulation error) (n = 10) or declined participation during the course of the experiment (e.g. pregnancy, claustrophobia) (n = 10). Sixty participants were included in the analysis (Table S1).

### Overall study design

This was a double-blinded, randomised, placebo-controlled cross-over study. For fMRI visits, participants arrived after an overnight fast and were provided with a standardized breakfast. After completing some computerized tasks which included training for in-scanner task on fMRI visit 1 and a working memory task battery on fMRI visit 2, they took either placebo or 150mg L-DOPA orally (counterbalanced across visits). This dosage was selected based on previous studies that have also used L-DOPA to investigate similar Markov decision making tasks [16,27]. About 45 minutes post-administration, during which L-DOPA plasma levels were expected to peak [28], participants completed the two-step task. The fMRI task took about 36 minutes. A washout period of at least 7 days between fMRI visits was implemented to prevent carry over treatment effects [29].

On the PET visit (mean interval ± s.d.: 78.2 ± 51.9 days post-fMRI visit 2), participants were administered 150mg Carbidopa orally 60 minutes prior to intravenous administration of ^18^F-DOPA. The PET scan took 4 hours. The experimental protocol was approved by the local ethics committee of Technische Universität Dresden (EK 44022012) and the German Federal Office for Radiation Protection (Bundesamt für Strahlenschutz). All participants provided written informed consent and received monetary compensation. Further details of the study procedure and data acquisition are reported in the Supplement.

### Behavioral data

Participants performed an adapted version of the two-step task [6] developed by Daw and colleagues [5] while undergoing fMRI (Figure S2). The rationale behind the task was to characterise participants based on their control strategies (MB or MF) during adaptive learning. The key feature that differentiated between these strategies was how first-stage choice behaviour depended on the ‘model’ i.e. task transition structure.

We conducted the behavioural analyses using a model-agnostic approach and a computational modelling approach. For the model-agnostic approach, we calculated individual scores for each control system to test how each component of the MB/MF balance is altered by DA. The probability to repeat the same first-stage choice in each upcoming trial depends on two conditions, namely the transition between first and second stage (common: C or rare: R), and the outcome at second-stage (rewarded: R or unrewarded: U) during the previous trial. A MB control user would tend to repeat (i.e. stay on) first-stage choices after common-rewarded (CR) trials and rare-unrewarded (RU) trials, whereas a MF control user would tend to repeat first-stage choices after being rewarded, disregarding the transition contingencies. As reported previously [30], MB and MF scores were calculated with probabilities to repeat i.e. stay on first-stage choices (P(stay)) for each type of trial as follows: MB score = P(stay|CR) - P(stay|RR) - P(stay|CU) + P(stay|RU), MF score = P(stay|CR) + P(stay|RR) - P(stay|CU) - P(stay|RU).

For comparison with findings from Deserno et al. [8], we employed the same computational approach to estimate the weighting parameter *ω* which indicates the balance between MF (*ω* = 0) and MB (*ω* = 1) control. In our previous work, which was based on a partially overlapping dataset with this current study, we found Daw’s original 7-parameter hybrid model to perform best in terms of model fit as compared to alternative models with fewer parameters [20]. In brief, the hybrid model was fitted to the behavioural data of each participant twice (please see Supplement for model specification). Data from both fMRI sessions were treated as a single group during the fitting procedure. Firstly, maximum likelihood was used to estimate the free parameters. Using these maximum likelihood estimates as prior distribution, the model was fitted again to the data using expectation maximisation to generate maximum-a-posteriori estimates. Bounded model parameters were transformed to an unconstrained scale via logistic transformation [*x*’=log/*x*(1-*x*))] for *α_1_, α_2_, λ, ω* and natural log transformation [*x*’=ln(*x*)] for *β_1_* and *β_2_* to adhere to normal distribution assumptions for model fitting and statistical analyses. The parameter estimates are shown in Table S3.

### Imaging data

Standard preprocessing procedure was applied to the data (please see detailed description in Supplement). First-level statistics were computed with a general linear model approach as reported before [5], using mean of individual model parameters to define the model, which was then used to generate individualised time series of RPEs by simulating the model over each individual’s experiences. MF learning signal was modelled with the regressor representing MF reward prediction error (MF-RPE). MB learning signal was modelled with the difference regressor (MB-dif), which represented the difference between MF-RPE and MB- RPE. Both MF-RPE and MB-dif were modelled as parametric modulators of second-stage and outcome onsets. These parameters were estimated on a voxel-wise basis before being averaged within three *a priori* regions-of-interest (ROI), namely ventral striatum (VST) and ventromedial prefrontal cortex (vmPFC) where Daw and colleagues have identified these learning signals [5] and lateral prefrontal cortex (LPFC) where Deserno et al. [8] have found an association between MB learning signal and DA synthesis capacity in the VST (Figure S3).

PET data were analysed as reported in our previous work [31]. In brief, we characterized three measures that reflected baseline DA, namely ^18^F-DOPA influx rate constant (*k_occ_*) which represented presynaptic DA synthesis capacity, ^18^F-DA washout rate (*k_loss_*) which reflected DA loss and effective distribution volume ratio (EDVR = *k_occ_/k_loss_*), which depicted ^18^F-DOPA trapped in the VST at *steady-state* [23,24]. *K_occ_* and EDVR were first estimated on a voxel-wise basis using a reversible-tracer graphical analysis approach [24,25] before being averaged within individualized VST ROIs. DA washout (*k_loss_*) was then calculated as the ratio *K_occ_* / EDVR for each ROI. As EDVR has been suggested to be a more sensitive indicator of baseline DA than *K_occ_* and *k_loss_* alone [25] and learning signals were associated with baseline DA within the VST, but not in caudate nor putamen [8], we used EDVR within the VST as our main index of baseline DA.

### Statistical analyses

Data were analyzed using MATLAB R2010 software (The Mathworks, Inc., MA, USA), IBM SPSS statistics 24 (IBM Corporation, NY, USA) and R Version 3.4.0 (https://www.r-project.org/). We assessed our main questions of interest, that is, whether there was a main effect of baseline DA, a main effect of L-DOPA intervention and, most importantly, an interaction between baseline DA and L-DOPA intervention on our two-step task measures reflecting MF and MB control. These measures included model-agnostic MF and MB scores, computational model estimate for *ω* and learning signals MF-RPE and MB-dif averaged within VST, vmPFC and lPFC. We assessed these effects for each task measure using separate repeated measures ANCOVAs, where the task measure of interest served as the dependent variable, drug condition as the independent variable and EDVR as a covariate.

As prior work showed practice effects in the two-step task [32], drug order and its interaction with drug condition was included to account for repetition effects of the cross-over design, and to factor out possible complex carry-over effects of treatment on practice effects e.g. participants tested with placebo first may gain a learning advantage during the L-DOPA session [33]. For assessing directionality for main effect of drug, we used paired *t*-test. For assessing directionality of the main effect of PET measures and their interactions with L-DOPA effects on two-step measures, we computed Pearson partial correlations, controlled for drug order. Main effects of PET measures are represented as correlations between PET measures and two-step measures averaged across both sessions. Interaction between PET measures and L-DOPA effects on two-step measures are represented as correlations between PET measures and L-DOPA induced change in two-step measures. Statistical significance for all analyses were based on *p* < 0.05, two-tailed, uncorrected for multiple comparisons.

## Results

### Participant characteristics

Our sample consisted of 60 healthy adults (mean age ± s.d.: 36.1 ± 3.80 years; mean body weight ± s.d.: 80.9 ± 12.6 kg; Table S1). They were predominantly male (n = 49), one-third were current smokers (n = 18) and drug order was fully counterbalanced. To check the validity of our measures of interest, we did as follows. We first assessed if our participants employed a mixture of MB and MF control during the task as reported previously [5,6,8]. Indeed, there was a main effect of reward and reward-by-transition interaction on first-stage stay probabilities (Table S2, Figure S4). We then assessed for MF and MB learning signals on a voxel-wise basis to visualise brain regions where BOLD signal during the second-stage and outcome onset corresponded with MF-RPE and MB-dif respectively (Figure S5). In keeping with previous work [5,6,8], strong MF-RPE signals were identified within the VST, whereas marginal MB-dif signals were identified within the vmPFC.

### DA modulation of MF and MB control behaviour

Main effects of EDVR, drug and their interaction on behavioural measures of MF and MB control are shown in Table 1. There was no main effect of EDVR on MF score and MB score (smallest *p* = .25). There was no main effect of drug on MB score (*p* = .16), but there was a main effect of drug on MF score (*F*(1,57) = 4.21, *p* = .045, *η_p_*^2^ = .069), where L-DOPA reduced the influence of previous-trial rewards on the tendency to stay on current first-stage choices (*t*(59) = -2.02; *p* = .048), indicative of reduced MF control. EDVR-by-drug interaction on MF score did not reach significance (*p* = .17). However, there was a EDVR-by-drug interaction on MB score (F(1,57) = 4.67, *p* = .035, *η_p_*^2^ = .076), where L-DOPA induced change in MB score, indicative of enhanced MB control, was greater in individuals with higher EDVR (*r* = 0.28, *p* = .035, Figure 1 top row). In order to clarify the drivers of this interaction, we conducted a *post-hoc* analysis where participants were median-split based on EDVR into two groups (median EDVR = 1.25). In the low-EDVR group, MB score did not differ (*p* = .81), whereas in the high-EDVR group, MB score was enhanced by L-DOPA (*t*(29) = 2.58, *p* = .015). As such, EDVR-by-drug interaction on MB score was driven by L-DOPA effects in high-EDVR individuals. As for the computational measure *ω* indicative of MB/MF balance, there were no main effects of EDVR, drug and their interaction on *ω* (smallest *p* = .06).

**Table 1.** Main effect of baseline DA, drug and their interaction on two-step behaviour.

| | MF score | | | MB score | | | $\omega$ | | |
| --- | --- | --- | --- | --- | --- | --- | --- | --- | --- |
| | F | Sig. | $\eta_p^2$ | F | Sig. | $\eta_p^2$ | F | Sig. | $\eta_p^2$ |
| EDVR | 1.38 | 0.246 | 0.024 | 0.25 | 0.619 | 0.004 | 0.78 | 0.381 | 0.013 |
| drug | **4.21** | **0.045** | **0.069** | 2.04 | 0.159 | 0.035 | 0.93 | 0.339 | 0.016 |
| EDVR-by-drug | 1.90 | 0.174 | 0.032 | **4.67** | **0.035** | **0.076** | 3.72 | 0.059 | 0.061 |
| drug order | 1.11 | 0.297 | 0.019 | 0.43 | 0.516 | 0.007 | 0.19 | 0.667 | 0.003 |
| drug order-by-drug | 1.47 | 0.231 | 0.025 | 2.97 | 0.090 | 0.049 | 0.25 | 0.616 | 0.004 |

In BOLD: $p < .05$

**Figure 1:**
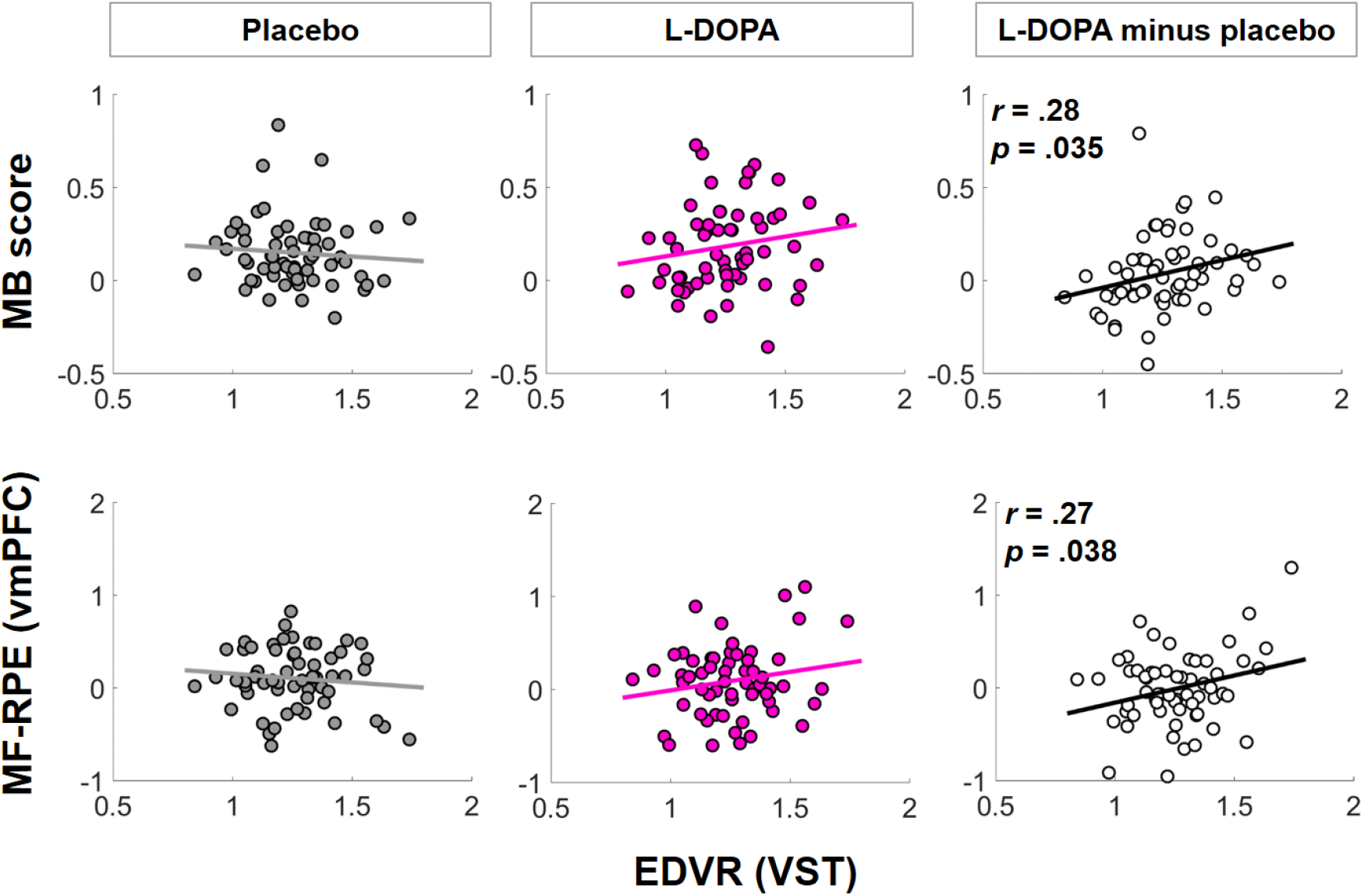
Individual differences in DA modulation of MB and MF control. Correlations between baseline DA (ventral striatal EDVR) and two-step behavioural and neural measures (for visualisation). (Top, left to right) MB score under placebo, L-DOPA and L-DOPA induced change in MB score. (Bottom, left to right) MF learning signal in the vmPFC under placebo, L-DOPA and L-DOPA induced change in MF learning signal.

### DA modulation of MF and MB learning signals

Main effects of EDVR, drug and their interaction of MF and MB learning signals are as shown in Table 2. There were no main effects of EDVR or drug on all learning signals (smallest *p* = .12). There was a significant EDVR-by-drug interaction on MF-RPE within the vmPFC (*F*(1,57) = 4.53, *p* = .038, *η_p_*^2^ = .074), where L-DOPA induced MF learning signal enhancement was greater in individuals with higher EDVR (*r* = 0.27, *p* = .038, Figure 1 bottom row). In order to clarify the drivers of this interaction, we conducted a *post-hoc* analysis as we did for the interaction on MB score. For both low and high-EDVR groups (divided based on a median split), MF-RPE did not differ significantly between drug conditions (smallest *p* = .54). EDVR-by-drug interaction on all other learning signals did not reach significance (smallest *p* = .06).

**Table 2.** Influence of higher DA levels (EDVR, L-DOPA) on MF and MB learning signals.

| MF-RPE | VST |  |  | vmPFC |  |  | IPFC |  |  |
| --- | --- | --- | --- | --- | --- | --- | --- | --- | --- |
| | F | Sig. | $\eta_p^2$ | F | Sig. | $\eta_p^2$ | F | Sig. | $\eta_p^2$ |
| EDVR | 0.29 | 0.593 | 0.005 | 0.234 | 0.631 | 0.004 | 2.43 | 0.124 | 0.041 |
| drug | 0.93 | 0.340 | 0.016 | 0.047 | 0.829 | 0.001 | 0.01 | 0.904 | 0.000 |
| EDVR-by-drug | 3.67 | 0.060 | 0.061 | **4.533** | **0.038** | **0.074** | 2.88 | 0.095 | 0.048 |
| drug order | 1.67 | 0.201 | 0.029 | 0.065 | 0.800 | 0.001 | 0.39 | 0.534 | 0.007 |
| drug order-by-drug | 0.47 | 0.494 | 0.008 | 2.311 | 0.134 | 0.039 | 0.02 | 0.885 | 0.000 |

| MB-dif | VST |  |  | vmPFC |  |  | IPFC |  |  |
| --- | --- | --- | --- | --- | --- | --- | --- | --- | --- |
| | F | Sig. | $\eta_p^2$ | F | Sig. | $\eta_p^2$ | F | Sig. | $\eta_p^2$ |
| EDVR | 0.79 | 0.379 | 0.014 | 0.85 | 0.361 | 0.015 | 0.53 | 0.468 | 0.009 |
| drug | 0.00 | 0.947 | 0.000 | 1.01 | 0.320 | 0.017 | 0.13 | 0.718 | 0.002 |
| EDVR-by-drug | 0.56 | 0.459 | 0.010 | 0.53 | 0.469 | 0.009 | 0.00 | 0.983 | 0.000 |
| drug order | 2.63 | 0.111 | 0.044 | 0.10 | 0.755 | 0.002 | 0.00 | 0.991 | 0.000 |
| drug order-by-drug | 3.84 | 0.055 | 0.063 | 0.33 | 0.565 | 0.006 | 0.00 | 0.987 | 0.000 |

In BOLD: $p < .05$

## Discussion

We investigated whether higher DA levels at baseline in the VST as measured with ^18^F-DOPA (a L-DOPA analogue) and L-DOPA induced enhancement in DA levels are associated with a shift in MB/MF balance towards MB control, and whether baseline DA interacts with the effect of L-DOPA on MB/MF balance. Baseline DA was not associated with MF control, MB control or MB/MF balance. L-DOPA reduced MF control on average across individuals, and enhanced MB control only for individuals with higher baseline DA. Our neural findings also showed a corresponding baseline-by-drug interaction, where higher-baseline DA individuals showed a greater L-DOPA induced enhancement in learning signals. Surprisingly, these observations were restricted to MF, rather than MB learning signals within the vmPFC.

Our main finding suggests that individuals with higher baseline DA showed greater L-DOPA induced enhancement in MB control. This is in concert with the general notion where higher DA levels is related to greater MB control. Of note, the interaction between resting striatal DA levels and L-DOPA effects on MB control were weak, albeit consistent in directionality across both model-agnostic and computational measures (Table S6a). As test-retest reliability of the VST has been shown to be poor when examined unilaterally [35], we chose to focus our main analyses on baseline DA averaged across both hemispheres. Notwithstanding, we found that these associations were weakened when analysed bilaterally, as these effects were restricted to the right VST (Table S7). One way to interpret our findings is that higher baseline DA individuals were better able to utilize L-DOPA as shown previously in rats [22]. As such, higher baseline DA individuals had a greater increase in DA levels under L-DOPA, thus increasing MB control. Upon further exploration of our data, we found that differential responses to L-DOPA we have observed in our study were driven by DA washout from the striatum rather than F-DOPA uptake (Table S6a, S7). High DA washout reflects a heightened DA turnover within the striatum, a process that can be influenced by different factors. Individuals with higher DA washout could be less efficient in storing DA and/or have faster DA catabolism mechanisms [36]. Based on animal studies that investigated the impact of DA storage and catabolism manipulation on L-DOPA changes in extracellular DA levels [37,38], we would expect individuals with high DA washout to have lower baseline DA and/or respond less to L-DOPA. Given that DA washout reflects multiple processes, future work should consider how each of these processes might interact with L-DOPA effects.

Deserno and colleagues found that individuals with higher DA synthesis capacity in the right VST showed a greater reliance on MB/MF balance, stronger lPFC MB learning signal and weaker VST MF learning signal. Wunderlich and colleagues showed that L-DOPA enhanced MB control. As we did not observe these associations in our data, we considered if this discrepancy might be related to model fit as our sample showed a poorer overall model fit than Deserno’s sample. When we excluded participants with poor model fit, our lack of findings with baseline DA remained unchanged, but we were able to replicate Wunderlich’s findings where L-DOPA enhanced MB control (N = 40; Table S8). With regard to the neural findings, we noted that the model parameters used for generating learning signals in Deserno’s study were treated as random effects, whereas we have treated them as mixed effects as was done in Daw’s study [5]. This may have reduced individual explanatory variance in our neural data, rendering it more challenging to identify associations with baseline DA [34]. In sum, although we did not observe an association between individual differences in baseline DA with MB and MF control as well as a L-DOPA enhancement on MB control across all participants, these effects may have been masked by model fit issues and/or analysis choice.

How do our findings further our understanding of the relationship between DA and MB control? A majority of previous work suggesting a positive relationship between baseline DA and MB control have related MB control to baseline executive function measures, such as WMC [10], which has been related to individual differences in baseline DA levels in the striatum [21] as well as the prefrontal cortex (PFC) [39]. In our sample, WMC showed a clear positive relationship with MB control averaged across sessions, consistent across both computational and model-agnostic measures (Table S9). Further, as pointed to previously [11], individuals with higher WMC also showed greater L-DOPA induced enhancement in MB control (Table S9). Although baseline striatal DA at rest was positively associated with WMC (*r* = .30, *p* = .018), resting striatal DA levels alone did not explain individual variability in MB control in our sample. Conventionally, WM performance has been related to an inverted U-shape relationship with D1 receptor activation in the PFC, where insufficient or excessive D1 activation can impair WM [40]. More recently, it has been suggested that an optimal WM performance relies on a balance between DA levels and D2 receptor availability within the PFC [41]. Future work could consider quantifying baseline DA levels as well as DA receptor binding within the PFC to gain a more comprehensive picture of how baseline DA functioning is related to MB control, and how DA modulatory effects on MB control depends on baseline DA functioning in the PFC.

What about MF control? Two studies found that individuals with higher ventral striatal DA synthesis capacity exhibited weaker MF learning signals [8,45]. Schlagenhauf *et al.* [45] suggested that higher striatal DA synthesis capacity reflects higher *tonic* DA levels, which is related to lower phasic DA-related neurotransmission manifested as reduced MF learning signal, whereas Deserno *et al.* [8] suggested that higher DA synthesis capacity is related to higher *phasic* DA transients, which manifested as greater uncertainty in MF estimates, causing a shift away from MF control. Although we did not observe negative associations between EDVR and MF control and MF learning signals (Table S6a,b), our exploratory analyses revealed that individuals with lower DA washout (with presumably higher baseline DA) showed less MF control. While this observation is supportive of the negative relationship between baseline DA and MF control, we exert caution in drawing further conclusions given its exploratory nature. MF learning relies on phasic DA firing of midbrain neurones [43], and its impact on choice behaviour is reduced when tonic DA levels are high [44]. Although it is unclear how resting baseline DA is related to tonic and phasic DA during the task, these findings seem to converge on the notion where higher baseline DA is related to a reduction in some aspect of MF control. In concert with this, we reported previously that enhancing DA levels with L-DOPA reduced MF control by attenuating the impact of learned reward information on future choice while leaving MF learning intact [20]. However, when we took into account individual differences in response to L-DOPA, we found that high-baseline individuals showed a L-DOPA induced enhancement in MF learning signals in the vmPFC, which is at odds with the ‘higher-DA-less-MF’ notion. Perhaps a way to reconcile these observations is to consider how L-DOPA might have enhanced tonic and phasic DA levels differentially across individuals. We would speculate that L-DOPA enhanced tonic DA levels similarly for all individuals, reducing MF control on average. However, L-DOPA enhancement of phasic DA levels may have been more dependent on baseline DA, thereby influencing MF learning signals. Future work should consider disentangling the influence of L-DOPA on phasic and tonic DA levels during the task, which should offer clarity on how DA is related to MF control.

There are several limitations in our study. We have reported previously that baseline DA is negatively associated with body mass index [48]. As such, it is possible that the interactions observed between baseline DA and L-DOPA effects reflect the influence of body weight on bioavailability, where individuals with higher baseline DA had a greater bioavailability of L-DOPA. Future studies should control for body weight when determining L-DOPA dosages for each participant. As we chose to restrict our fMRI analyses to RPE signals for consistency with prior work [5,8], we did not investigate other neural processes previously implicated in mediating MB/MF balance. The inferior lateral prefrontal cortex has been shown to confer a bias towards MB control by altering connectivity between prefrontal and striatal regions [49]. Extending our work to connectivity measures could shed further light on the mechanisms underlying our observations with MB control, which we could not explain by observing MB learning signals alone. Given that our PET measure has only been shown to provide reliable information about DA functioning within the striatum [35], we were unable to explore the role of baseline DA in extra-striatal regions. Lastly, due to our recruitment strategy [50], our sample was biased in terms of gender ratio and 5-HTTLPR genotype. As such, generalisability of our results is limited.

Taken together, we have sought to clarify the relationship between baseline DA, L-DOPA effects and their interaction on MB and MF control during reinforcement learning. To our knowledge, this was the first study of its kind that characterised the role of DA in MB and MF control using a baseline measure reflecting striatal DA levels at rest and a complementary pharmacological intervention during the task. Overall, our results support previous work pointing to the relationship between higher DA levels and greater MB control. Individual differences in baseline DA within the ventral striatum account for differential effects of L-DOPA on MB control and MF learning signals.

## Acknowledgements

MNS and TG designed the study. NBK was responsible for running the study, YL coordinated and ran the data collection. YL performed the data analysis, with contributions from NBK, MNS, SP, LO, LD, QJMH and DM. YL wrote the first draft of the manuscript. All authors contributed and have approved the final manuscript. We thank the SFB940/1 Project B3 and B4 study teams, as well as colleagues at the Neuroimaging Center and Department of Nuclear Medicine for providing technical support.

