## Supplemental Material for "Individual differences in dopamine function underlying the balance between model-based and model-free control"

### Supplement

#### Study procedure

*Recruitment*

We sent invitation letters to individuals randomly selected by the residents' registration office of Dresden, Germany (N = 15778) to recruit a community sample. Participants were included for the study if they (1) were between 30 to 40 years old at the time of recruitment, (2) had no history of neurological or mental disorders according to the Screening Version of the Structured Clinical Interview for DSM-IV (Wittchen *et al*, 1997) except for nicotine dependence, (3) had no MRI, PET nor L-DOPA contraindications, (4) were physically able to lie comfortably in the scanner for at least 4 hours, (5) had normal or corrected-to-normal vision, (6) had no current use of illicit drugs nor alcohol consumption.

During the planning stages of this study, there were no appropriate data available for a formal effect size calculation. Given that fMRI signal-to-noise ratio is typically low (Desmond and Glover, 2002), and prior studies that investigated baseline DA relationships with related constructs (e.g. working memory capacity) reported effect sizes of $r = .45$ (Landau et al., 2009) and $\rho = .68$ (Cools et al., 2008), we expected to detect at least medium-sized effects (i.e. $r = 0.3$) (Cohen, 1988). To maintain Type 1 error probability at 0.05 and power at 0.8, we targeted for 60 participants which would be sufficient for detecting effects with $r > 0.35$, two-tailed (Faul *et al*, 2007).

*Experimental visits*

All participants attended four experimental visits: a baseline visit and two functional magnetic resonance imaging (fMRI) visits at the Neuroimaging Center at the Technische Universität Dresden, followed by a positron emission tomography (PET) visit at the PET-Center (Department of Nuclear Medicine, Technische Universität Dresden).

During the baseline visit, blood samples were drawn for genotyping for a partner study (Neukam *et al*, 2018) along with height and weight measurements before participants completed a series of computerized and pen-and-paper questionnaires.

Each participant then attended two fMRI visits. In order to minimize the influence of psychoactive drugs on BOLD signal and food on levodopa absorption (Crevoisier *et al*, 2003), participants were asked to abstain from medication for at least 24 hours prior to their visit and arrive at the scanning facility after an overnight fast. Checks for current use of illicit drugs (urine test on first fMRI visit only; Kombi/DOA10-Schnelltest, MAHSAN Diagnostika GmbH, Reinbek, Germany) and alcohol consumption (breath-alcohol test on both fMRI visits;

Alcotest 6510, Drägerwerk AG & Co. KGaA, Lübeck, Germany) were then administered. Participants were given a small standardised breakfast (total: ~25g butter biscuits, ~120 kcal) upon arrival (between 05:30 – 08:45 hrs) and dextrose tablets throughout the session (total: ~4g, 17,4 kcal per hour) to reduce side effects of the drug administration. Participants completed a series of computerised tasks before being administered orally 150mg/37.5mg L-DOPA/benserazide (Madopar, 125 mg T, Roche, Switzerland) or placebo (P-Tabletten, 8mm Lichtenstein Winthrop Arzneimittel GmbH, Germany). The computerised tasks included training for the in-scanner task (for first fMRI) and operation span task (for second fMRI). The order of drug condition during the fMRI visits was pseudo-randomised across participants. Participants then entered the MRI scanner for structural scans before they began with the in-scanner task.

On the day of PET visit, participants were asked to abstain from protein-containing foods. When the participants arrived at the PET-centre (10am), 150mg Carbidopa (Amerigen Pharmaceuticals Inc., NJ, USA) was administered orally to increase $^{18}$F-DOPA bioavailability (i.e. plasma levels) and thereby maximising cerebral uptake of $^{18}$F-DOPA (Hoffman *et al*, 1992). Weight was measured on the day of PET visit, while height was measured during baseline. BMI was calculated as kg/m$^2$. One hour post-Carbidopa administration, a mean $^{18}$F-DOPA activity of 172 ± 8.43 (range 120 – 185) MBq was administered intravenously at the start of the PET scan. The PET scan followed a 4-hour acquisition protocol developed by Sossi and colleagues (Sossi *et al*, 2002). All participants were instructed beforehand to keep still during the scan.

*MRI acquisition*

MRI images were acquired on a 3-Tesla Magnetom Trio Tim system (Siemens, Erlangen, Germany) with a 32-channel head coil. During the in-scanner task, stimuli were presented via an MR compatible screen and rearview mirror system. Participants responded by pressing their index fingers on two separate button boxes, one held in each hand. Psychophysics Toolbox (Version 3) (Brainard, 1997; Kleiner *et al*, 2007; Pelli, 1997) implemented within MATLAB R2010a software (The Mathworks, Inc., MA, USA) was used for stimulus presentation and behavioural data collection. Functional images were acquired using a gradient echo-planar imaging (EPI) sequence, repetition time TR = 2.41 s; echo time TE = 25 ms; flip angle: 80°; field of view: 192 x 192 mm$^2$; matrix size: 64 x 64; voxel size: 3 x 3 x 2 mm$^3$ (slice thickness: 2 mm; gap: 1 mm). Every volume consisted of 42 transverse slices acquired descending from the top, manually adjusted ~25° clockwise from the anterior commissure-posterior commissure plane (total ~900 volumes for each participant, total scan time: ~36min). A corresponding field map was also recorded for distortion correction of the EPI images. Structural images were acquired using a T1-weighted magnetization prepared rapid acquisition with gradient echo (MPRAGE) sequence for normalization, anatomical localization as well as screening for structural abnormalities by a neuro-radiologist (TR: 1.90 s; TE: 2.52 ms; flip angle: 9°; field of view: 256 x 256 $mm^2$; number of volumes: 192; voxel size: 1 x 1 x 1 $mm^3$).

*PET acquisition*

PET images were acquired with an Ingenuity TF PET/MR scanner (3T; Philips Healthcare, OH, USA). 3D emission data were acquired in list-mode. The acquisition comprised three phases (phase 1: 0-min post-injection for 90-min, phase 2: 130-min post-injection for 40-min, phase 3: 200-min post-injection for 40-min), where every phase was preceded by an MR scan for attenuation correction. Participants were able to take break(s) outside of the scanner between phases. A 4-min structural MR T1 image was also acquired using a T1-weighted (TFE) sequence with a SENSE 8-channel head coil for anatomical localization (TR: 8 ms; TE: 4 ms; flip angle: 8°; field of view: 240 x 240 $mm^2$; number of slices: 192; voxel size: 1.0 x 1.0 x 1.0 $mm^3$).

*Two-step task*

The original two-step task (Daw *et al*, 2011) was adapted for the following: (i) instructions were translated into German, (ii) visual stimuli were adapted to present different sets of stimuli across visits (pseudo-randomised across participants), and (iii) outcome presentation times at both stages were decreased by a factor of 2 to reduce trial duration. The task consisted of a total of 201 trials, separated by inter-trial intervals sampled from an exponential distribution (mean = 2 s, range: 1-7 s). Each trial consisted of two stages (Figure S2A). At the first stage, the participants had to choose between two grey stimuli. After the first-stage choice, they were led to a second stage where they had to make a choice between two coloured (green/yellow) stimuli. After they have made the second-stage choice, monetary outcome (win 20 cents or 0 cents) for the trial was presented. Over time, participants had to learn two aspects of the task: (i) the transition structure (Figure S2B), which grey stimulus led to the yellow pair of stimuli at 70% of the trials ('common trials') and to the green pair at 30% of the trials ('rare trials') (and vice versa for the other grey stimulus); and (ii) the reward probabilities associated with each second-stage stimulus, which followed Gaussian random walks that changed slowly and independently of each other with reflecting boundaries at 0.25 and 0.75 (Figure S2C). Prior to the experiment, participants were explicitly instructed on these two aspects of the task. They were asked to choose the best option (i.e. with highest reward probability) on every trial during the task. At the end of the fMRI visit 2, we explicitly asked the participants which strategies they have employed during the two-step task on both fMRI visits. As this post-fMRI interview was implemented shortly after data collection has begun, we only had this information for a subset of our participants (n = 50). They were classified as having employed a MB strategy for both visits if they took the transition probabilities into consideration when making their first-stage choices (n = 31).

*Operation span task*

The operation span task was part of a working memory task battery (Lewandowsky *et al*, 2010) and it was administered between a memory updating task and a spatial short-term memory test. For each trial of the operation span task, participants were presented with a series of alternating arithmetic equations and consonants. Length of each trial (i.e. list length) ranged between 4 to 8 pairs of equations and letters. When an equation (e.g. '3 + 3 = 5') was presented on the screen, the participant had to press the left button if the equation was correct, and the right button if it was false. At the end of the trial, participants had to type the consonants in the order they were presented. There were 15 trials in total, 3 trials per list length, and took about 10min in total. The task was slightly modified from the original version. Practice trials were changed from three trials without feedback to two trials with feedback and included an option to be repeated. For the actual task, letter sequences, equations and trial order that were previously identical for all participants was modified such that the sequence of list length was kept identical for all participants, but the letters and equations presented were randomised across participants.

Working memory capacity (WMC) was calculated as follows. The task had two components, namely letters to be remembered and arithmetic equations be processed. The proportion of letters remembered in the correct order was first averaged across all trials. Similarly, the proportion of equations processed correctly was also averaged across all trials. Instead of having two separate outcome measures as depicted in the original paper (Lewandowsky *et al*, 2010), we combined them into a single score (OS score) by converting both subscores into standardised z-scores before summing them together.

#### Behavioural data analyses

Behavioural data were preprocessed using MATLAB R2010 software (The Mathworks, Inc., MA, USA) and IBM SPSS statistics (IBM Corporation, NY, USA).

*Computational modeling*

The task consisted of three states, $s_A$ at first stage, $s_B$ and $s_C$ at second stage. At each state, the participant has to decide which action ($a_A$ and $a_B$) to take in order to maximize his/her rewards from the task. Based on reinforcement learning theory, the participant decides which action to take by predicting how good it is to perform each action in a given state (Sutton and Barto, 1998). In other words, the participant learns the expected future value for each state-action pair $Q$ ($s$, $a$) during the task. Here, we assume that the participant learns these values by employing a hybrid of two learning methods: namely MF and MB learning. It has been shown previously that this hybrid model fits best to participants' behaviour when compared to other simpler variants of this model (Deserno *et al*, 2015; Sebold *et al*, 2017).

This section describes the MF learning algorithm. At stage $i$ of trial $t$, the participant computes $Q_{MF}$ of the next trial as follows:

$$Q_{MF}(s_{i,t+1}, a_{i,t+1}) = Q_{MF}(s_{i,t}, a_{i,t}) + \alpha_i \delta_{i,t} \text{ [S1]}$$

where

$$\delta_{i,t} = r_{i,t} + Q_{MF}(s_{i+1,t}, a_{i+1,t}) - Q_{MF}(s_{i,t}, a_{i,t}). \text{ [S2]}$$

MF reward prediction error (MF-RPE) i.e. MF learning signal was modelled with $\delta$, which represents the difference between actual outcome and expected value (Note: at first stage, reward $r_{1,t} = 0$; at second stage, $Q_{MF}(s_3,a_{3,t}) = 0$). $\alpha_1$ and $\alpha_2$ are the learning rates at first and second stages respectively, that is, the extent $Q_{MF}$ is updated by current information $\delta$. When an error occurs at the end of the trial, eligible state-action pairs are assigned credit for the error (Sutton *et al*, 1998) as depicted here:

$$Q_{MF}(s_{1,t+1}, a_{1,t+1}) = Q_{MF}(s_{1,t}, a_{1,t}) + \alpha_1 \lambda \delta_{2,t}, \text{ [S3]}$$

where $\delta$ at second-stage is weighted (i.e. given more credit) by eligibility trace $\lambda$ for updating $Q_{MF}$ at first-stage of the next trial. This MF algorithm is also known as Sarsa(λ) (Sutton *et al*, 1998).

This section describes the MB learning algorithm. Here, for each first-stage action, the participant first estimates the value attributed to each second-stage state by multiplying its transition probability (i.e. the chance of arriving at this second-stage state given the first-stage action) with its corresponding $Q_{MF}$ of the better action. $Q_{MB}$, the expected value of each first-stage action, is then given by the sum of these two values as shown here:

$$Q_{MB}(s_A, a_j) = P(s_B|s_A, a_j) \max_a Q_{MF}(s_B, a) + P(s_C|s_A, a_j) \max_a Q_{MF}(s_C, a), \text{ [S4]}$$

where $a \in \{a_A, a_B\}$ , $P\ (s_B|s_A,a_A) = 0.7$, $P\ (s_C|s_A,a_B) = 0.7$ or $P\ (s_B|s_A,a_A) = 0.3$, $P\ (s_C|s_A,a_B) = 0.3$. (Note: at second-stage, $Q_{MB} = Q_{MF}$.) By learning this transition function which maps the state-action values to the transition probabilities, the participant takes the task structure into consideration (hence 'model-based').

Finally, this section describes the hybrid learning algorithm, which combines both MF and MB learning into the calculation of net state-action value $Q_{net}$ at first-stage:

$$Q_{net}(s_A, a_j) = \omega Q_{MB}(s_A, a_j) + (1 - \omega) Q_{MF}(s_A, a_j), \text{ [S5]}$$

where $\omega$ indicates the relative influence of MF and MB learning on $Q_{net}$. Net RPE is calculated as follows:

$$\delta_{net,i,t} = r_{i,t} + Q_{net}(s_{i+1,t}, a_{i+1,t}) - Q_{net}(s_{i,t}, a_{i,t}). \text{ [S6]}$$

MB learning signal was modelled with the difference regressor (MB-dif), which was calculated as the difference between $\delta_{net}$ when $\omega = 1$ and $\omega = 0$.

Action probabilities are then calculated as a function of expected values of state-action pairs based on softmax action selection rule:

$$P(a_{i,t} = a|s_{i,t}) = \frac{\exp(\beta_i[Q_{net}(s_{i,t},a) + \pi \cdot \mathrm{rep}(a)])}{\sum_{a'}\exp(\beta_i[Q_{net}(s_{i,t},a') + \pi \cdot \mathrm{rep}(a')])}, \text{ [S7]}$$

where $\beta_1$ and $\beta_2$ determine choice consistency (also known as inverse temperature, where a high $\beta$ indicates high correspondence between action probabilities and their expected values) at first and second stage respectively, rep($a$) depicts whether $a$ was the same as (=1) or differed from (=0) the one chosen on the previous trial and $\pi$ represents the degree of perseveration on first-stage choices.

#### Functional data analyses

fMRI data were analysed with SPM8 (Wellcome Trust Centre for Neuroimaging, London, UK) implemented within Nipype Version 0.9.2 (Gorgolewski *et al*, 2011) and MATLAB R2010 software (The Mathworks, Inc., MA, USA).

*Preprocessing*

fMRI data were preprocessed as follows. The first 4 volumes of the EPI images were discarded to allow for magnetic saturation. The remaining 896 volumes were subjected to slice time correction (reference: middle slice), followed by realignment to the first volume of the run to correct for motion. Distortion correction based on the field map was then applied to the realigned EPI images. Each individual anatomical T1 image was first co-registered to the individual mean EPI image before segmentation and normalization to MNI space. The resulting transformation parameters were then applied to the distortion-corrected EPI images to spatially normalize them to MNI space (resampled to the final voxel size: 2 x 2 x 2 $mm^3$). Finally, normalized EPI images were spatially smoothed with an isotropic Gaussian kernel (full width at half maximum: 8 mm). During first-level analyses, the data was high-pass-filtered at 128 s.

*MF-RPE and MB-dif*

This analysis is based on the computational modelling approach as presented by Daw et al. (Daw *et al*, 2011). For the first-level analyses of each participant, fMRI signal was modelled in a voxel-wise manner using a general linear model described as follows. The first regressor corresponded to onset of second stage stimulus and outcome presentation, followed by two parametric regressors which were second stage and outcome onset weighted by MF-RPE (Equation S2) and MB-dif (Equation S6). Regressors of non-interest included one regressor for outcome onset, one regressor for first stage stimulus onset, followed by two parametric regressors which were first stage stimulus onset weighted by probabilities of first stage actions (Equation S7) and its partial derivative with respect to ω, one regressor for missed trials and six regressors (three translation and three rotation parameters) for capturing residual movement-related artifacts. Each regressor was convolved with a canonical hemodynamic response function. L-DOPA and placebo conditions were modelled separately. Parameter estimates for MF-RPE and MB-dif were then taken to a second-level random-effects model.

#### Potentially confounding variables

A confounder is defined as a variable which alters the association(s) between independent and dependent variables of interest (Greenland and Robins, 2009). Variables of interest included all measures from the two-step task, including behavioural ($\omega$, MB score, MF score in placebo/L-DOPA condition) and neural (MF-RPE, MB-dif averaged in VST, vmPFC, LPFC) measures, and all PET measures (EDVR, $k_{occ}$, $k_{loss}$ averaged in VST). Sex and smoking status were included for consideration as potentially confounding variables as they have been associated with dopamine synthesis capacity (Laakso *et al*, 2002; Rademacher *et al*, 2016). Body weight was also included for consideration as we found that individuals with higher body-mass index had lower striatal EDVR (Lee *et al*, 2018). Further, body weight has been shown to influence pharmacokinetics of L-DOPA (Zappia *et al*, 2002) and a dose-dependent effect on cognition due to body weight has been shown in a previous study that used the same L-DOPA dosage as in our study (Rutledge *et al*, 2015). Each of these potentially confounding variables were assessed individually for associations with variables of interest using separate one-way MANOVAs (see Table S4) where all variables of interest were entered as dependent variables, and each potential confound was entered as a between-subject factor (categorical) or a covariate (continuous).

There were no significant differences in variables of interest between male and female groups (smallest $p$ = .06) nor smoker groups (smallest $p$ = .06). Higher body weight was associated with lower EDVR ($F(1,59)$ = 7.13, $p$ = .010, $r$ = - .33) and higher $k_{loss}$ ($F(1,59)$ = 5.15, $p$ = .027, $r$ = .29) in the VST. Since sex and smoking status were not significantly associated with any variables of interest, only body weight was included as potential confounds. When we repeated our main behavioural analysis with body weight included, we found that body weight did not interact significantly with drug effects on each behavioural measure (Table S5). Given that body weight did not seem to influence bioavailability of L-DOPA, we did not include body weight as a confound since it would reduce variance of interest.

**Table S1. Participant characteristics**

| Variables of interest | Mean ± S.D. (Min – Max) | | Ratio |
|---|---|---|---|
| **Age (years)** | 36.1 ± 3.80 (30.0 – 42.1) | **Sex (male/female)** | 49/11 |
| **Weight (kg)** | 80.9 ± 12.6 (52.6 – 113.6) | **Smoking status (current-/ex-/non-smokers)** | 18/17/25 |
| | | **Drug order (L-DOPA first/placebo first)** | 30/30 |
| ***Baseline DA*** | | ***MB/MF balance ω***[a] | |
| **EDVR** | 1.26 ± 0.18 (0.84 – 1.74) | **Placebo** | 0.41 ± 0.27 (0.02 – 0.97) |
| $k_{occ}$ | 0.0153 ± 0.0014 (0.0113 – 0.0176) | **L-DOPA** | 0.49 ± 0.32 (0.01 – 0.99) |
| $k_{loss}$ | 0.0139 ± 0.0018 (0.0105 – 0.0182) | | |

[a]Values shown here are bounded between 0 and 1 for interpretability

**Table S2. Effects on first-stage stay probabilities**

| Effect of interest | F | Sig. | $\eta_p^2$ |
|---|---|---|---|
| drug | 0.31 | 0.581 | 0.01 |
| drug * drug order | 0.63 | 0.430 | 0.01 |
| reward | **41.07** | **<.001** | **0.41** |
| reward * drug order | 1.15 | 0.287 | 0.02 |
| transition | **9.98** | **0.003** | **0.15** |
| transition * drug order | 2.78 | 0.101 | 0.05 |
| drug * reward | **4.10** | **0.048** | **0.07** |

| | | | |
|---|---|---|---|
| drug * reward * drug order | 1.38 | 0.245 | 0.02 |
| drug * transition | 2.56 | 0.115 | 0.04 |
| drug * transition * drug order | 1.45 | 0.233 | 0.02 |
| reward * transition | **48.48** | **<.001** | **0.46** |
| reward * transition * drug order | 0.42 | 0.519 | 0.01 |
| drug * reward * transition | 1.86 | 0.178 | 0.03 |
| drug * reward * transition * drug order | 2.66 | 0.109 | 0.04 |
| drug order | 0.43 | 0.516 | 0.007 |

Stay probability: probability of repeating first-stage choice
Drug: L-DOPA/placebo
Drug order: L-DOPA first/placebo first
Reward: rewarded/unrewarded on previous trial
Transition: common/rare transition on previous trial

**Table S3. Distribution of best-fitting parameters and negative log-likelihood**

| | | $\beta_1$ | $\beta_2$ | $\alpha_1$ | $\alpha_2$ | $\lambda$ | $\omega$ | $\pi$ | nLL |
|---|---|---|---|---|---|---|---|---|---|
| **Placebo** | 25th percentile | 2.55 | 1.56 | 0.31 | 0.37 | 0.33 | 0.18 | 0.12 | 241.72 |
| | 50th percentile | 3.93 | 2.60 | 0.54 | 0.56 | 0.67 | 0.41 | 0.18 | 201.37 |
| | 75th percentile | 7.75 | 4.11 | 0.79 | 0.79 | 0.95 | 0.66 | 0.30 | 169.82 |
| **L-DOPA** | 25th percentile | 3.38 | 1.21 | 0.26 | 0.35 | 0.23 | 0.19 | 0.06 | 240.65 |
| | 50th percentile | 5.30 | 2.41 | 0.45 | 0.57 | 0.67 | 0.57 | 0.14 | 204.88 |
| | 75th percentile | 8.33 | 3.74 | 0.84 | 0.76 | 0.94 | 0.77 | 0.23 | 166.52 |

$\alpha_1$ and $\alpha_2$: first and second stage learning rates; $\beta_1$ and $\beta_2$: first and second stage choice consistencies; $\lambda$: eligibility trace; $\pi$: perseveration; $\omega$: relative influence of MF and MB; nLL: negative log-likelihood

**Table S4. Potential confounds**

| | | Gender | | | Smoker Status | | | Weight | | | |
|---|---|---|---|---|---|---|---|---|---|---|---|
| | | F | Sig. | $\eta_p^2$ | F | Sig. | $\eta_p^2$ | F | Sig. | $\eta_p^2$ | $r$ |
| **PET VST** | **EDVR** | 0.267 | 0.608 | 0.005 | 0.251 | 0.779 | 0.009 | **7.132** | **0.010** | **0.110** | **-0.33** |
| | $k_{occ}$ | 2.447 | 0.123 | 0.040 | 0.187 | 0.830 | 0.007 | 3.779 | 0.057 | 0.061 | -0.25 |
| | $k_{loss}$ | 0.583 | 0.448 | 0.010 | 1.043 | 0.359 | 0.035 | **5.146** | **0.027** | **0.081** | **0.29** |
| **Behaviour** | $\omega^L$ | 0.182 | 0.671 | 0.003 | 1.834 | 0.169 | 0.060 | 0.170 | 0.681 | 0.003 | -0.05 |
| | $\omega^P$ | 0.043 | 0.837 | 0.001 | 0.441 | 0.646 | 0.015 | 0.788 | 0.378 | 0.013 | 0.12 |
| | **MF score**$^L$ | 0.008 | 0.927 | 0.000 | 2.395 | 0.100 | 0.078 | 0.529 | 0.470 | 0.009 | 0.10 |
| | **MF score**$^P$ | 1.293 | 0.260 | 0.022 | 0.970 | 0.385 | 0.033 | 0.232 | 0.632 | 0.004 | -0.06 |
| | **MB score**$^L$ | 0.541 | 0.465 | 0.009 | 1.927 | 0.155 | 0.063 | 0.200 | 0.656 | 0.003 | -0.06 |
| | **MB score**$^P$ | 0.090 | 0.765 | 0.002 | 1.005 | 0.373 | 0.034 | 2.259 | 0.138 | 0.037 | 0.19 |
| **fMRI VST** | **MF-RPE**$^L$ | 1.382 | 0.245 | 0.023 | 3.057 | 0.055 | 0.097 | 0.677 | 0.414 | 0.012 | -0.11 |
| | **MF-RPE**$^P$ | 0.020 | 0.889 | 0.000 | 1.123 | 0.332 | 0.038 | 1.441 | 0.235 | 0.024 | 0.16 |
| | **MB-dif**$^L$ | 0.068 | 0.796 | 0.001 | 1.988 | 0.146 | 0.065 | 1.920 | 0.171 | 0.032 | -0.18 |
| | **MB-dif** $^P$ | 0.071 | 0.791 | 0.001 | 1.361 | 0.265 | 0.046 | 0.001 | 0.979 | 0.000 | 0.00 |
| **vmPFC** | **MF-RPE**$^L$ | 0.250 | 0.619 | 0.004 | 0.717 | 0.493 | 0.025 | 0.003 | 0.957 | 0.000 | 0.01 |
| | **MF-RPE**$^P$ | 0.367 | 0.547 | 0.006 | 1.351 | 0.267 | 0.045 | 1.717 | 0.195 | 0.029 | 0.17 |
| | **MB-dif**$^L$ | 0.001 | 0.979 | 0.000 | 0.783 | 0.462 | 0.027 | 0.755 | 0.389 | 0.013 | -0.11 |
| | **MB-dif** $^P$ | 3.726 | 0.058 | 0.060 | 1.001 | 0.374 | 0.034 | 0.597 | 0.443 | 0.010 | -0.10 |
| **lPFC** | **MF-RPE**$^L$ | 0.190 | 0.664 | 0.003 | 2.342 | 0.105 | 0.076 | 0.095 | 0.759 | 0.002 | 0.04 |
| | **MF-RPE**$^P$ | 0.408 | 0.526 | 0.007 | 0.535 | 0.588 | 0.018 | 1.284 | 0.262 | 0.022 | 0.15 |

| | | | | | | | | | |
|---|---|---|---|---|---|---|---|---|---|
| **MB-dif**$^{L}$ | 0.084 | 0.774 | 0.001 | 3.071 | 0.054 | 0.097 | 0.314 | 0.578 | 0.005 | -0.07 |
| **MB-dif** $^{P}$ | 0.323 | 0.572 | 0.006 | 0.527 | 0.593 | 0.018 | 0.120 | 0.730 | 0.002 | -0.05 |

L: L-DOPA session, P: placebo session

**Table S5. Main effect of baseline DA, drug and their interaction on two-step behaviour, including body weight**

| | $\omega$ | | | **MB score** | | | **MF score** | | |
|---|---|---|---|---|---|---|---|---|---|
| | **F** | **Sig.** | $\eta_p^2$ | **F** | **Sig.** | $\eta_p^2$ | **F** | **Sig.** | $\eta_p^2$ |
| EDVR | 0.98 | 0.325 | 0.017 | 2.04 | 0.159 | 0.035 | 1.33 | 0.253 | 0.023 |
| drug | 0.91 | 0.343 | 0.016 | 2.72 | 0.105 | 0.046 | **4.22** | **0.045** | **0.070** |
| EDVR-by-drug | 2.87 | 0.096 | 0.049 | 1.41 | 0.240 | 0.025 | 0.88 | 0.353 | 0.015 |
| drug order | 0.16 | 0.694 | 0.003 | 2.71 | 0.105 | 0.046 | 1.06 | 0.307 | 0.019 |
| drug order-by-drug | 0.27 | 0.604 | 0.005 | 0.49 | 0.488 | 0.009 | 1.29 | 0.260 | 0.023 |
| weight | 0.25 | 0.621 | 0.004 | 0.48 | 0.493 | 0.008 | 0.03 | 0.862 | 0.001 |
| weight-by-drug | 0.11 | 0.742 | 0.002 | 0.48 | 0.489 | 0.009 | 1.21 | 0.276 | 0.021 |

In BOLD: $p < .05$

**Table S6a. Relationships between PET measures with two-step behaviour**

| | | $\mu$ | | | $\delta$ | | |
|---|---|---|---|---|---|---|---|
| | | **MF score** | **MB score** | $\omega$ | **MF score** | **MB score** | $\omega$ |
| **EDVR** | $r$ | -0.153 | 0.066 | 0.116 | -0.180 | **0.275** | 0.247 |
| | **Sig.** | .246 | .619 | .381 | .174 | **.035** | .059 |
| $k_{occ}$ | $r$ | 0.022 | -0.019 | -0.121 | -0.169 | 0.119 | 0.214 |
| | **Sig.** | .867 | .885 | .363 | .200 | .371 | .104 |
| $k_{loss}$ | $r$ | **0.261** | -0.141 | -0.229 | 0.133 | **-0.288** | -0.193 |
| | **Sig.** | **.046** | .287 | .081 | .314 | **.027** | .142 |

*r*: Pearson's partial correlation, controlled for drug order
In BOLD: $p$ <.05
$\mu$: averaged across sessions
$\delta$: L-DOPA minus placebo session

**Table S6b. Relationships between PET measures with MF and MB learning signals**

| **MB-dif** | | *μ* | | | *δ* | | |
|---|---|---|---|---|---|---|---|
| | | **VST** | **vmPFC** | **lPFC** | **VST** | **vmPFC** | **lPFC** |
| **EDVR** | ***r*** | -0.117 | -0.121 | -0.096 | 0.098 | 0.096 | -0.003 |
| | **Sig.** | .379 | .361 | .468 | .459 | .469 | .983 |
| $k_{occ}$ | ***r*** | 0.033 | 0.034 | -0.046 | 0.081 | 0.103 | 0.090 |
| | **Sig.** | .804 | .801 | .729 | .541 | .439 | .500 |
| $k_{loss}$ | ***r*** | 0.099 | 0.125 | 0.051 | -0.086 | -0.055 | 0.049 |
| | **Sig.** | .456 | .346 | .699 | .520 | .679 | .714 |

| **MF-RPE** | | *μ* | | | *δ* | | |
|---|---|---|---|---|---|---|---|
| | | **VST** | **vmPFC** | **lPFC** | **VST** | **vmPFC** | **lPFC** |
| **EDVR** | ***r*** | -0.071 | 0.064 | 0.202 | 0.246 | **0.271** | 0.219 |
| | **Sig.** | .593 | .631 | .124 | .060 | **.038** | .095 |
| $k_{occ}$ | ***r*** | -0.003 | 0.065 | 0.049 | -0.033 | 0.086 | 0.033 |
| | **Sig.** | .983 | .622 | .714 | .806 | .517 | .802 |
| $k_{loss}$ | ***r*** | 0.047 | -0.017 | -0.168 | **-0.314** | -0.197 | -0.170 |
| | **Sig.** | .726 | .899 | .205 | **.015** | .134 | .198 |

*r*: Pearson's partial correlation, controlled for drug order
In BOLD: $p$ <.05
*μ*: averaged across sessions
*δ*: L-DOPA minus placebo session

**Table S7. Relationships between PET measures with two-step behaviour (right versus left ventral striatum)**

| **Right** | | $\delta$ | | | **Left** | $\delta$ | | |
|---|---|---|---|---|---|---|---|---|
| | | **MF score** | **MB score** | $\omega$ | | **MF score** | **MB score** | $\omega$ |
| **EDVR** | ***r*** | -0.206 | **0.378** | **0.341** | **EDVR** | -0.120 | 0.134 | 0.118 |
| | **Sig.** | .118 | **.003** | **.008** | | .364 | .313 | .372 |
| $k_{occ}$ | ***r*** | -0.152 | 0.117 | 0.215 | $k_{occ}$ | -0.133 | 0.077 | 0.136 |
| | **Sig.** | .252 | .378 | .101 | | .317 | .563 | .304 |
| $k_{loss}$ | ***r*** | 0.138 | **-0.375** | **-0.311** | $k_{loss}$ | 0.080 | -0.117 | -0.033 |
| | **Sig.** | .299 | **.003** | **.016** | | .548 | .379 | .805 |

*r*: Pearson's partial correlation, controlled for drug order
In BOLD: $p$ <.05
$\delta$: L-DOPA minus placebo session

**Table S8. Main effect of baseline DA, drug and their interaction on two-step behaviour (good model-fit subgroup)**

| | **MF score** | | | **MB score** | | | $\omega$ | | |
|---|---|---|---|---|---|---|---|---|---|
| | **F** | **Sig.** | $\eta_p^2$ | **F** | **Sig.** | $\eta_p^2$ | **F** | **Sig.** | $\eta_p^2$ |
| EDVR | 1.08 | 0.306 | 0.03 | 0.42 | 0.521 | 0.01 | 1.00 | 0.323 | 0.03 |
| drug | 3.49 | 0.070 | 0.09 | **5.86** | **0.021** | **0.14** | 1.64 | 0.208 | 0.04 |
| EDVR-by-drug | 1.89 | 0.177 | 0.05 | **4.73** | **0.036** | **0.11** | 1.11 | 0.300 | 0.03 |
| drug order | 0.41 | 0.526 | 0.01 | 0.72 | 0.402 | 0.02 | 0.02 | 0.893 | 0.00 |
| drug order-by-drug | 0.89 | 0.352 | 0.02 | **5.33** | **0.027** | **0.13** | 1.14 | 0.292 | 0.03 |

In BOLD: $p$ <.05; good model-fit subgroup (N=40): bottom 75% in negative log-likelihood (nLL) for both sessions. High nLL (as in highly negative) reflects how poorly the estimated model predicts the individual's choice behaviour.

**Table S9. Relationships between working memory capacity with EDVR in the VST, two-step behavioural measures on average and their alterations with L-DOPA**

| | | | ***μ*** | | | ***δ*** | | |
|---|---|---|---|---|---|---|---|---|
| | | **EDVR** | **MF score** | **MB score** | ***ω*** | **MF score** | **MB score** | ***ω*** |
| **WMC** | ***r*** | **0.304** | -0.112 | **0.331** | **0.282** | -0.079 | **0.256** | **0.358** |
| | **Sig.** | **0.018** | 0.399 | **0.01** | **0.03** | 0.55 | **0.05** | **0.005** |

*r*: Pearson's partial correlation, controlled for drug order
WMC: working memory capacity measured using operation span
In BOLD: $p$ <.05

**Figure S1. Flow diagram of participant recruitment**

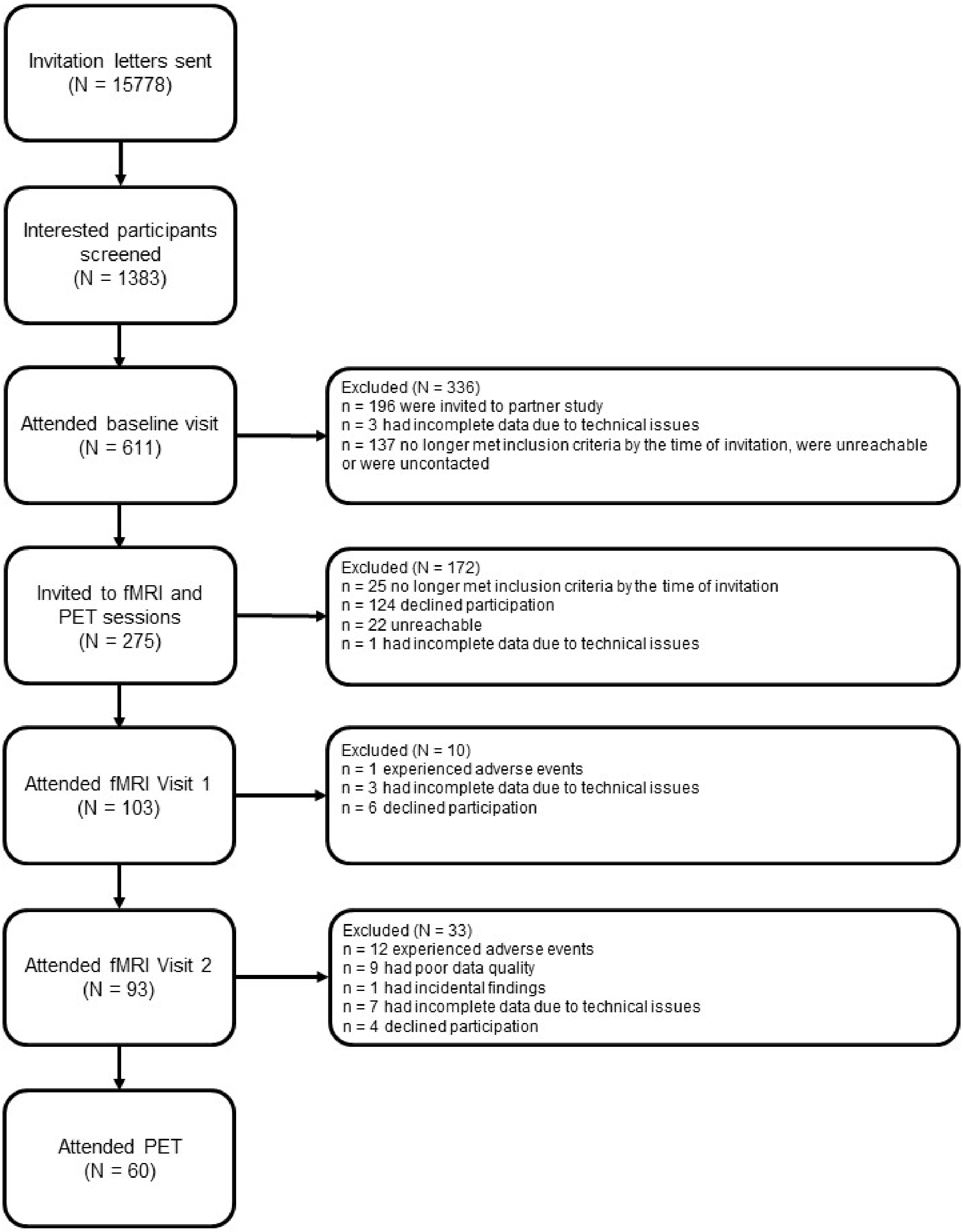

#### Figure S2: Schematic view of the two-step task

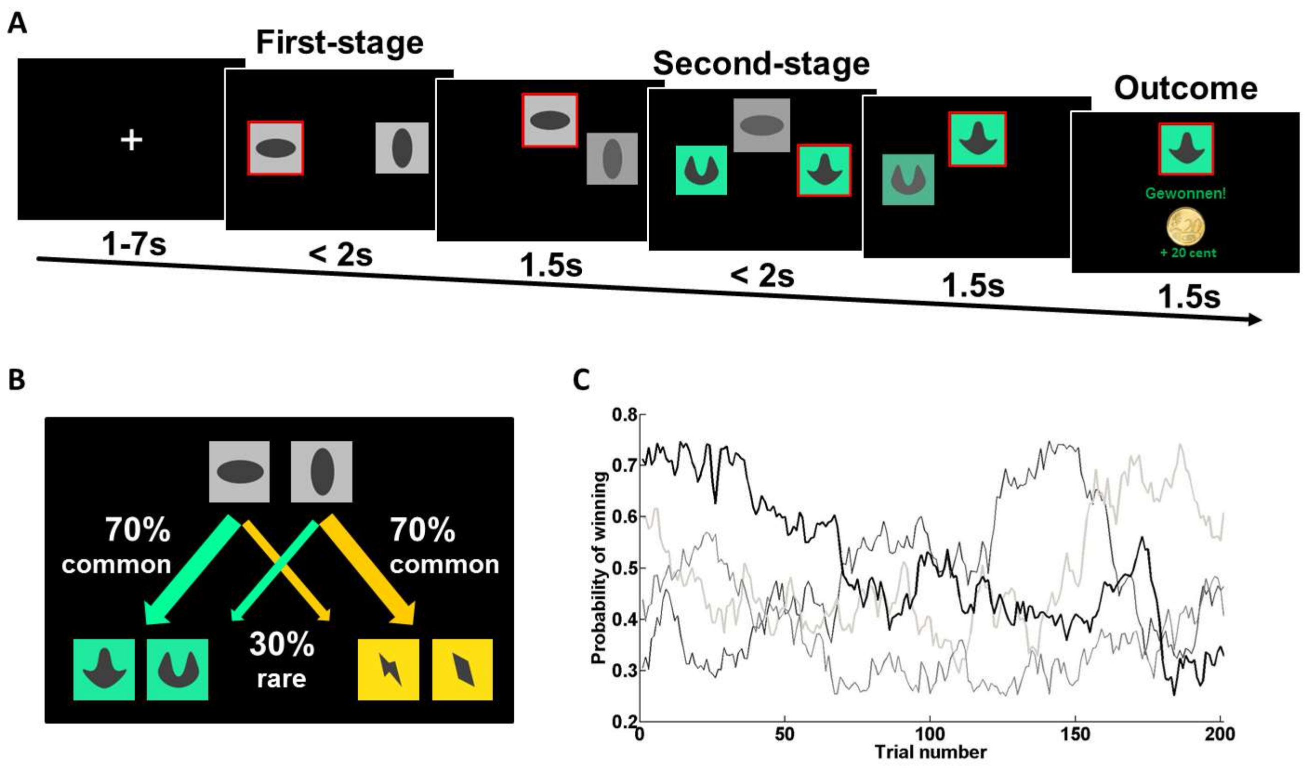


Figure S2: Schematic view of the two-stage Markov decision task (Figure adapted from (Sebold *et al*, 2017)). A: Trial sequence; B: Structure of transition contingencies; C: Reward probabilities for each of the four second-stage options across the whole experiment.

#### Figure S3. Regions of interest

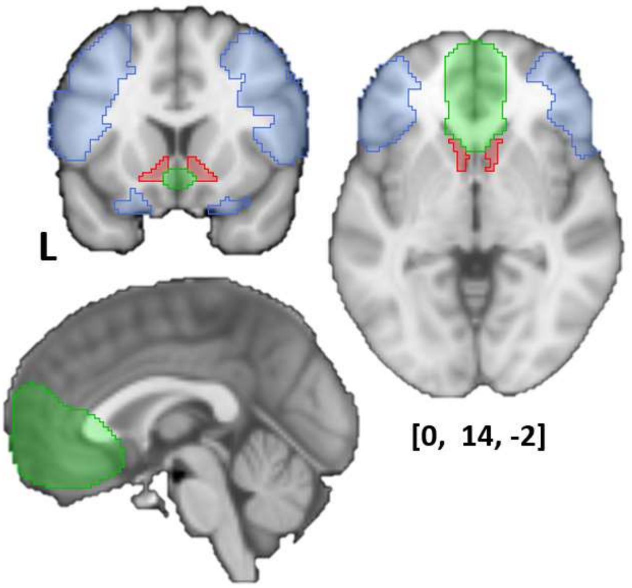


Figure S3: Regions of interest in MNI space. Ventral striatum (VST; red) were defined as reported by Tziortzi and colleagues (Tziortzi *et al*, 2011). vmPFC (green) was defined using Neurosynth as described by Wagner for the liberal version (Wagner, 2016) followed by exclusion of voxels which overlapped with the striatum. lPFC (blue) was defined as reported by Deserno and colleagues, , comprising the middle and inferior frontal gyrus of the Automated Anatomic Labeling Atlas (Deserno *et al*, 2015).

#### Figure S4. Stay probabilities of first-stage choices

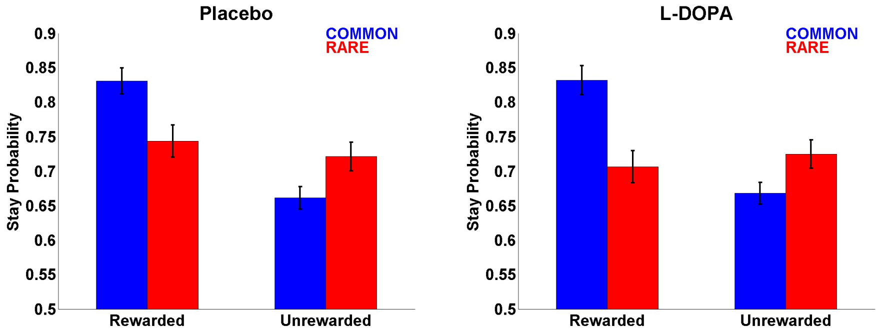


Figure S4: Stay probability of first-stage choices for current-trials separated by previous-trial conditions under placebo (left) and L-DOPA (right). There was a significant main effect of reward and reward-by-transition interaction (Table S2), which indicates that participants employed a mixture of MF and MB control strategies during the task.

#### Figure S5: Whole-brain MF and MB learning signals

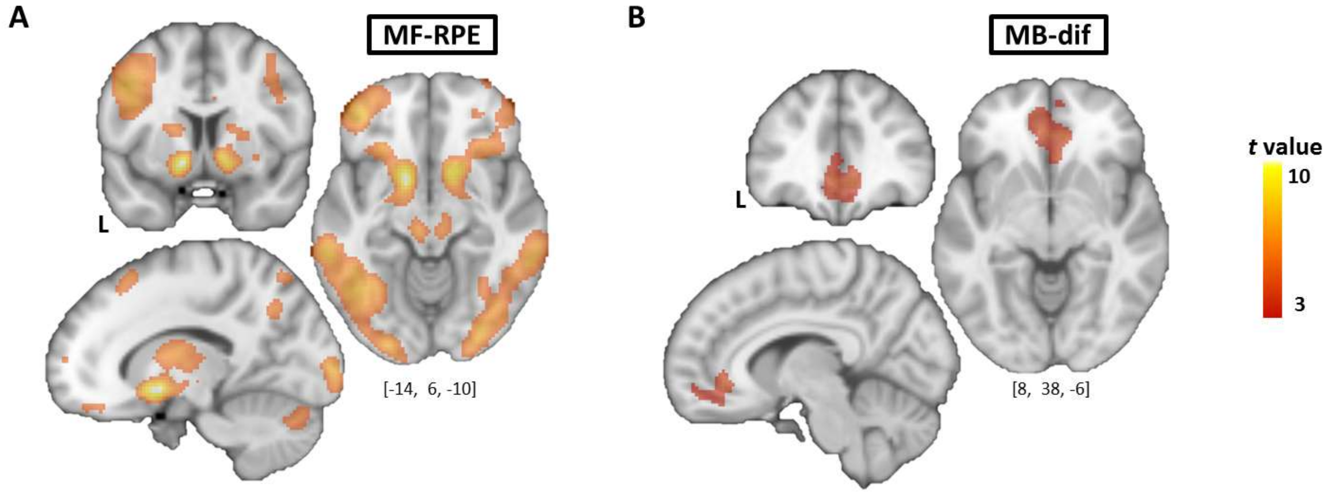


Figure S5: Whole-brain BOLD associations with MF and MB learning. A: Statistical map for main effect of MF-RPE (FWE corrected at $p < .05$, peak at left nucleus accumbens (ventral striatum): -14, 6, -10); B: Statistical map for main effect of MB-dif ($p < .001$ uncorrected, cluster extent k=20, peak at right cingulate: 8, 38, -6); regions were labelled according to the ICBM atlas implemented within WFU PickAtlasTool
